# Plasma Proteomics of Genetic Brain Arteriosclerosis and Dementia Syndrome Identifies Signatures of Fibrosis, Angiogenesis, and Metabolic Alterations

**DOI:** 10.1101/2024.03.28.587249

**Authors:** Jonah N. Keller, Hannah Radabaugh, Nikolaos Karvelas, Stephen Fitzsimons, Scott Treiman, Maria F. Palafox, Lisa McDonnell, Yakeel T. Quiroz, Francisco J. Lopera, Debarag Banerjee, Michael M. Wang, Joseph F. Arboleda-Velasquez, James F. Meschia, Adam R. Ferguson, Fanny M. Elahi

## Abstract

Cerebral autosomal dominant arteriopathy with subcortical infarcts and leukoencephalopathy (CADASIL) is the most common monogenic form of vascular cognitive impairment and dementia. A genetic arteriolosclerotic disease, the molecular mechanisms driving vascular brain degeneration and decline remain unclear. With the goal of driving discovery of disease-relevant biological perturbations in CADASIL, we used machine learning approaches to extract proteomic disease signatures from large-scale proteomics generated from plasma collected from three distinct cohorts in US and Colombia: CADASIL-Early (*N* = 53), CADASIL-Late (*N* = 45), and CADASIL-Colombia (*N* = 71). We extracted molecular signatures with high predictive value for early and late-stage CADASIL and performed robust cross- and external-validation. We examined the biological and clinical relevance of our findings through pathway enrichment analysis and testing of associations with clinical outcomes. Our study represents a model for unbiased discovery of molecular signatures and disease biomarkers, combining non-invasive plasma proteomics with clinical data. We report on novel disease-associated molecular signatures for CADASIL, derived from the accessible plasma proteome, with relevance to vascular cognitive impairment and dementia.

## Introduction

Cerebral autosomal dominant arteriopathy with subcortical infarcts and leukoencephalopathy (CADASIL) is an autosomal dominant form of VCID caused by missense mutations in *NOTCH3*. CADASIL is the leading cause of hereditary stroke and vascular cognitive impairment and dementia. Although the classical Mendelian syndrome is considered rare, with a prevalence of 1.3 - 4.1 per 100,000 adults (*1*), variants in *NOTCH3* associated with endophenotype of white matter disease are more common, occurring in as many as 1 in 300 individuals (*2*). Such prevalence suggests that discoveries of mechanisms underlying VCID in CADASIL could be relevant to vascular white matter disorders and dementia syndromes.

Research on the molecular pathogenesis of CADASIL has largely been limited to mice and postmortem human brain tissue. Consequently, our understanding of the early and evolving molecular pathogenesis of CADASIL, most relevant for development of impactful therapeutics, remains limited (*3*). NOTCH3 is a transmembrane receptor that is highly expressed in mural cells. A key unresolved question involves understanding how mutations in the extracellular domain (ECD) of NOTCH3 lead to the dysfunction of small-caliber blood vessels (*4*) and multicellular vascular phenotypes such as neurovascular decoupling, hypoperfusion, and blood-brain barrier dysfunction (*5*). In humans, neuroimaging abnormalities provide indications of affected vascular microenvironments (*6–8*). As a chronic disease, molecular pathologies in CADASIL unfold over decades. Although NOTCH3 is predominantly expressed by brain vascular mural cells, which molecular dysfunctions drive clinical symptomatology and how to counter disease progression across various disease stages is not understood. A comprehensive and unbiased molecular investigation could capture key molecular drivers of brain dysfunction and degeneration in CADASIL, highlighting potential therapeutic opportunities.

In this study, we generated unbiased plasma proteomics and asked whether CADASIL has a specific disease signature in peripheral blood in early and late stages of disease and whether these signatures are associated with clinically relevant outcomes. We used multivariate analytical methods, such as machine learning (ML), to identify molecular signatures of disease. Leveraging publicly available brain single cell transcriptomics, we assigned proteomic signatures to cells within the neurovascular unit. Finally, using clinical outcomes of relevance to VCID, we demonstrated the clinical relevance of our findings, with implications for future novel biomarker development for risk stratification, prognostication, and disease monitoring across spectrum of disease severity from early to late stage of CADASIL.

Here, we investigated the plasma proteome of CADASIL using data generated from an aptamer-based assay that quantifies over 7,000 proteins (SomaSCAN 7k, Somalogic, Boulder, CO) (*9–12*). Three distinct CADASIL cohorts with different disease stages were included in our analyses. Considering the age-associated nature of CADASIL, in which cognitive impairment and disability typically manifest after the age of 55 years (with notable patient-to-patient variability), we categorized our cohorts accordingly (*13, 14*). The older cohort from the Mayo Clinic (*N* = 45; *M_Age_* > 55 years) was labeled as CADASIL-Late, while the younger cohort recruited at UCSF (*N* = 53; *M_Age_* < 55 years) was designated as CADASIL-Early. We employed the third cohort from Colombia, South America as a holdout validation dataset (CADASIL-Colombia; *N* = 71), owing to technical variations in its collection and processing. To identify proteomic signatures intrinsically linked to the disease, we developed a novel machine learning methodology. Our methodological workflow incorporates consensus aggregation of a suite of statistical evaluators coupled with rigorous cross-validation. Our overarching objectives were to: (1) isolate early- and late-stage CADASIL proteomic signatures; (2) validate these signatures both internally and in external CADASIL populations; (3) elucidate the biological implications of these identified proteins; and (4) correlate these protein signatures with relevant clinical and imaging metrics.

The goal of our study was to uncover disease-associated molecular signatures with a hypothesis-agnostic approach to allow for data-driven discovery of molecular perturbations and identification of novel therapeutic targets. To this end, we leveraged state-of-the-art computational methods to analyze plasma proteome data generated in three distinct CADASIL cohorts spanning early preclinical disease to more advanced clinical stages involving strokes. In addition, we used brain tissue to validate the expression of key proteins and associated molecular pathways in brain tissue donated from individuals with CADASIL.

## Results

An overview of the study design is presented in **Figure 1**. The demographic characteristics of each cohort are described in **Table 1**. Designation of CADASIL-Early and CADASIL-Late monikers was based on the mean age of CADASIL participants and their symptom severity in respective cohorts.

**Fig. 1.**
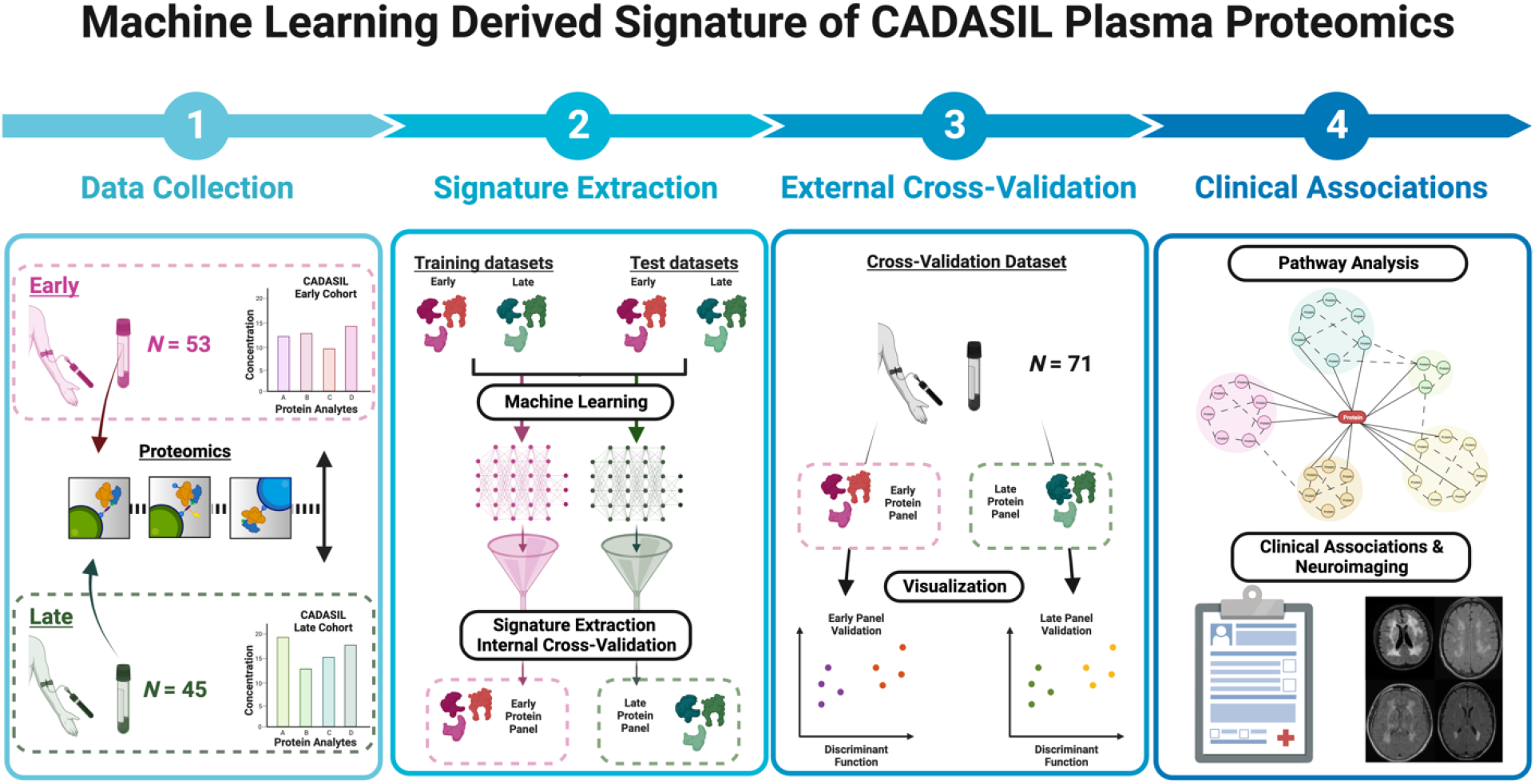
Illustration of data collection and analytical workflow used in this study. Plasma samples from three distinct CADASIL cohorts were collected and analyzed using the SomaSCAN proteomics platform. Protein signatures were determined by applying a machine learning feature extraction methodology, and findings were validated both internally and externally, as well as on post-mortem brain tissue. Protein panels were interrogated using interactive enrichment analysis, and we investigated the associations of molecular measures with quantitative measures of clinical abnormalities.

**Table 1.** Summary Demographic Data of CADASIL Cohorts. Control, healthy control; CADASIL, CADASIL cases.

| Cohort | Status | Sample Size (N) | Female (%) | Age (yrs.)<br>(mean ± SD) |
| --- | --- | --- | --- | --- |
| Early | Control | 28 | 50 | 52 (± 11) |
| | CADASIL | 25 | 36 | 51 (± 12) |
| Late | Control | 25 | 20 | 64 (± 11) |
| | CADASIL | 20 | 50 | 57 (± 12) |
| Colombia | Control | 42 | 40 | 41 (± 15) |
| | CADASIL | 29 | 28 | 39 (± 10) |

### Novel ML workflow detects CADASIL-associated proteomics signatures in peripheral blood

To elucidate the proteomic signatures of CADASIL, we developed a hypothesis-agnostic, multivariate analytical workflow, emphasizing robust statistical validation and biological and clinical interpretation of findings (**Fig. 2**). Our initial challenge pertained to reducing the dimensionality of over 7,000 proteins to more manageable protein lists. To achieve this, we implemented the leave-one-out (LOO) method, creating partitions (LOO folds) of our dataset. This method ensured minimal bias from individual samples and enhanced the generalizability of our model. In each LOO fold, only proteins with significant differences across groups (*P* < 0.05) progressed to subsequent analyses, resulting in approximately 1,300 proteins per fold. We then used a diverse array of feature selection algorithms, with different mathematical decision boundaries and solvers, carefully chosen to address the unique challenges posed by the high dimensionality and low sample size of our dataset. Next, we adopted a novel highly stringent, heterogeneous ensemble aggregation technique, which combines various algorithmic predictions to achieve a more accurate and reliable outcome. This method was crucial in ensuring that our protein selection was not biased towards any single machine learning algorithm, nor overfitted to any one sample. Proteins were only included in our final proteomic signature if they passed the highly conservative requirement of selection across all LOO folds by at least two different evaluators. When applied to each discovery cohort, these criteria led to the identification of two definitive protein signatures: the CADASIL-Early signature with 16 unique proteins and the CADASIL-Late signature with 20 unique proteins.

**Fig. 2.**
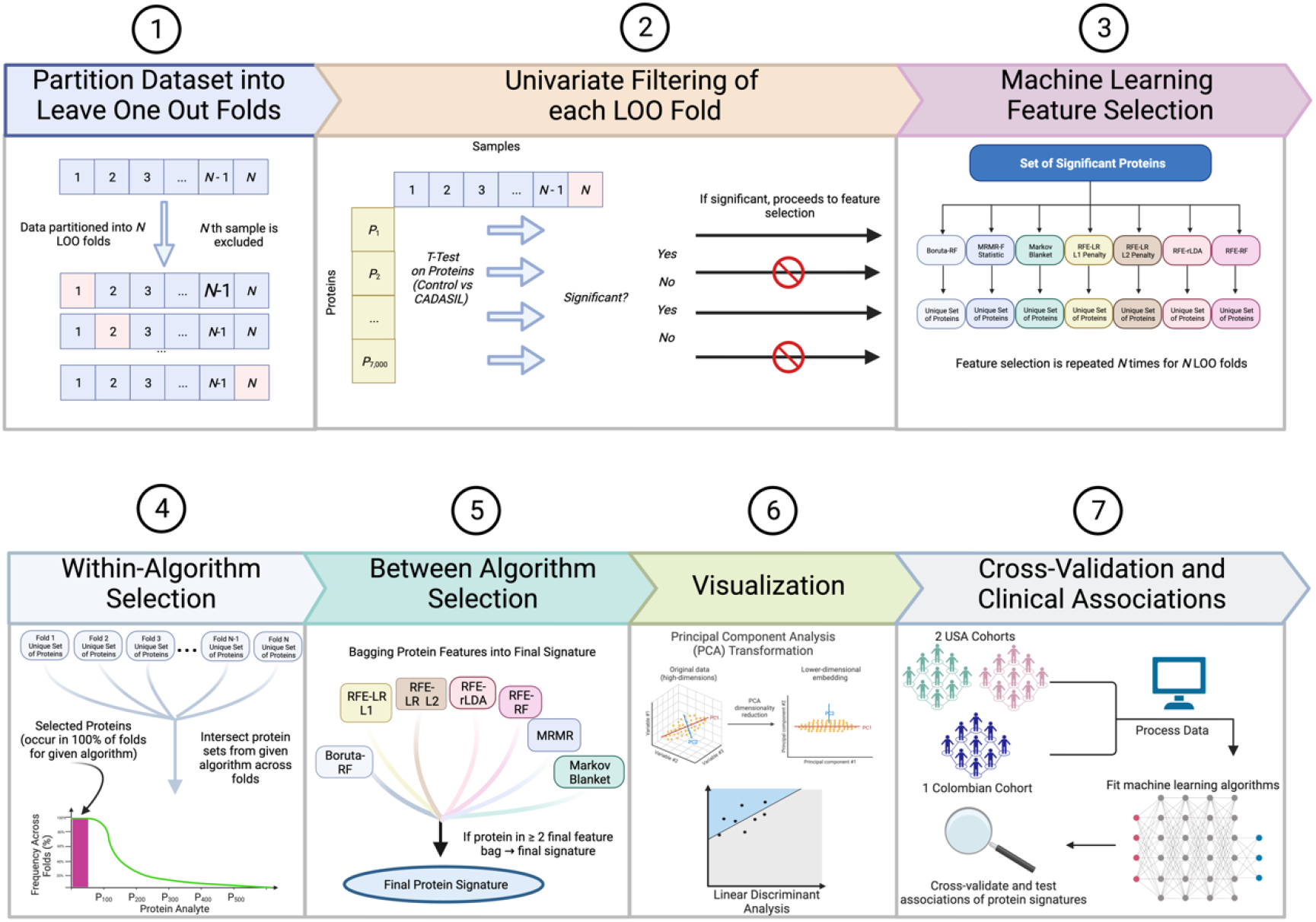
Illustration of machine learning methodology. Multivariate analytical workflow for CADASIL proteomic signature identification, featuring robust statistical validation, dimensionality reduction from 7,000 to 1,300 proteins, LOO method for bias minimization, diverse feature selection algorithms, and a stringent ensemble aggregation technique leading to two definitive protein signatures.

The following 16 proteins composed the CADASIL-Early proteomic signature: ANP32B, C4A|C4B, ENPP2, FN1, FUT3, GAS7, GPX1, HMBS, HPCAL1, MB, MGP, MYZAP, RRM1, SPINK6, UROS, and VEGFR3 (**Fig. 3A**). The following 20 proteins composed the CADASIL-Late proteomic signature: ABO, ACAA1, B3GAT1, BCAR3, C4A|C4B, CD209, DTNA, ECH1, ENDOU, FABP4, HNMT, KLRF1, KNG1, LTA4H, MASP1, OMG, SEMA3B, SLITRK1, TARDBP, and TMEM132B (**Fig. 3B**).

**Fig. 3.**
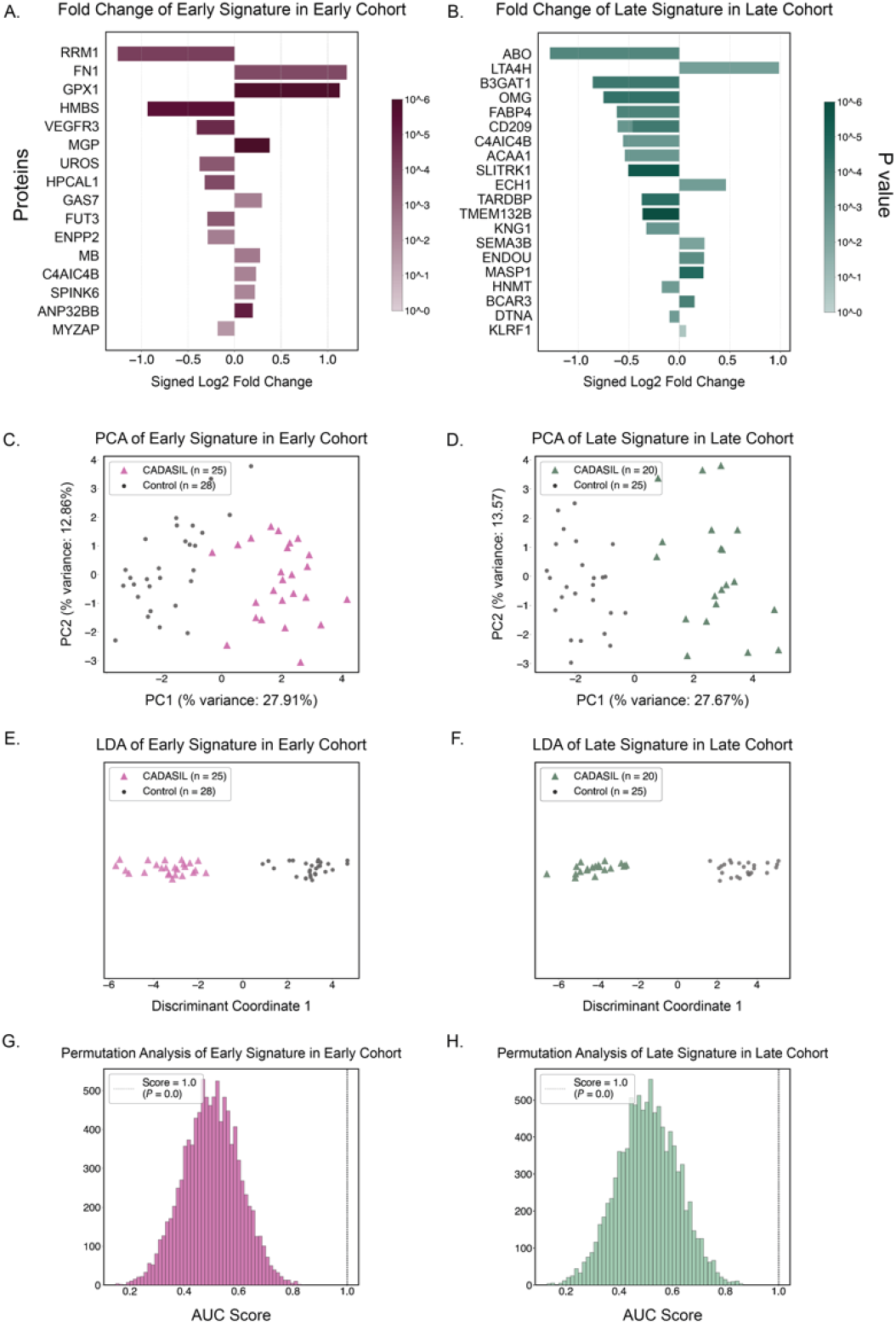
Machine learning identifies highly accurate and precise plasma proteomic signatures that discriminate CADASIL from control groups in early and late disease stages. **(A)** Fold changes (log_2_FC) of Early signature protein expression between CADASIL and Control groups. **(B)** Fold changes (log_2_FC) of Late signature protein expression between CADASIL and Control groups. **(C)** Principal component analysis of CADASIL-Early cohort separated from Control group, using Early signature proteins as features. **(D)** Principal component analysis of CADASIL-Late cohort using Late signature proteins as features. **(E)** Regularized linear discriminant analysis of CADASIL-Early cohort using Early signature proteins as features. **(F)** Regularized linear discriminant analysis of CADASIL-Late cohort using Late signature proteins as features. **(G)** Histogram of permutation results of random feature selection performance on Early cohort compared to derived Early signature. **(H)** Histogram of permutation analysis on Late cohort compared to derived Late signature.

### Machine learning model differentiates individuals affected with CADASIL from Healthy Controls using protein signatures

To further validate the uncovered protein signatures, we visualized associations with disease states using both unsupervised and supervised machine learning approaches. Principal component analysis (PCA), a non-supervised multivariate method, was performed on the selected set of proteins included in the Early and Late signatures (16 and 20 proteins, respectively). Visualization of the data on the principal component axes highlighted the elimination of non-disease-related signals (**Fig. 3C-D**), following the curation steps. We then trained supervised regularized linear discriminant analysis (rLDA) models using the protein sets as the input and disease status as the output. This approach resulted in ideal classification of disease instances (**Fig. 3E-F**). Rigorous cross-validation of the rLDA models was performed, which showcased their capacity for high accuracy and precision in distinguishing CADASIL from control samples, with statistical significance surpassing that of the permuted (i.e. random protein) models (*P* < 0.00001; **Fig. 3G-H**).

### Internal and external validation of the CADASIL-Late protein signature for distinguishing CADASIL patients from Controls

To assess the generalizability and reproducibility of our protein signatures, we performed both an internal validation using the opposing cohort (i.e., Late vs Early) in addition to an external validation using an independently collected dataset shared by the Neuroscience Group of Antioquia (Colombia) with data generated using plasma samples collected from a Colombian cohort (CADASIL-Colombia; demographics in **Table 1**). In the internal validation label permutation testing, we found that the CADASIL-Early signature was noisy in discriminating CADASIL from control groups in the CADASIL-Late cohort (*P* > 0.05; **Fig. 4A**). However, the CADASIL-Late signature performed at highly significant levels when distinguishing between CADASIL and control subjects in the CADASIL-Early cohort (*P* = 0.004; **Fig. 4B**).

**Fig. 4.**
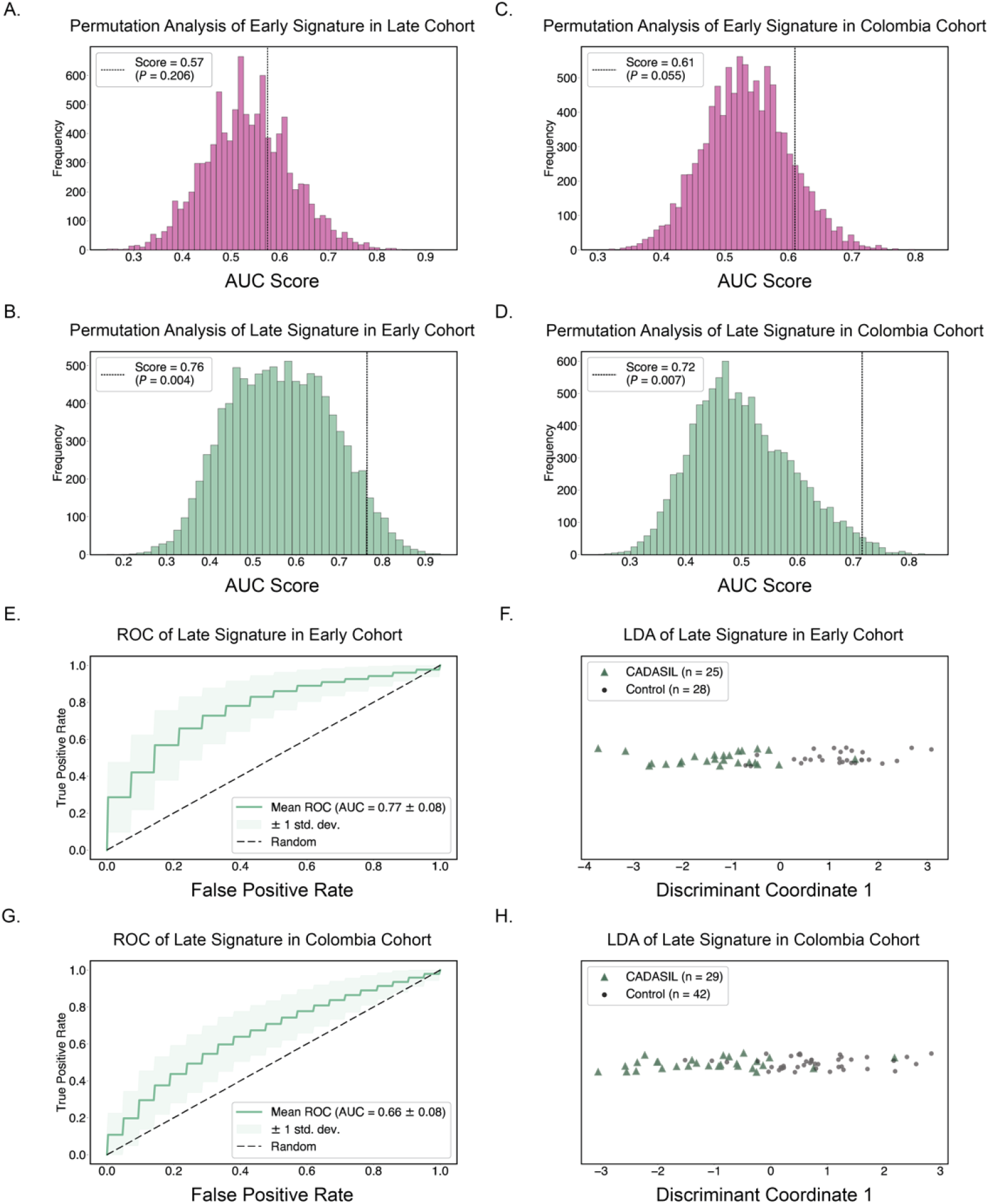
Validation of machine learning-derived Early and Late signatures. **(A)** Permutation analysis of Early signature performance in Late Cohort. **(B)** Permutation analysis of Late signature performance in Early Cohort. **(C)** Permutation analysis of Early signature in the external Colombia Cohort. **(D)** Permutation analysis of Late signature in the external Colombia Cohort. **(E, F)** Biplot combining ROC curve and LDA plot for Late signature in Early Cohort. **(G, H)** Biplot of ROC curve and LDA plot for Late signature in Colombia Cohort.

In our external validation, we trained supervised classifiers to distinguish disease status of the CADASIL-Colombia cohort using the Early and Late protein signatures as input. The CADASIL-Early plasma signature was marginally significant in discriminating in the CADASIL-Colombia dataset (*P* = 0.055; **Fig. 4C**). However, the CADASIL-Late plasma signature was significant in discriminating in the CADASIL-Colombia dataset (*P* = 6.6 × 10^−3^; **Fig. 4D**). To further substantiate these findings, we conducted permutation-based testing on the CADASIL-Late results and investigated separation in the high dimensional space by plotting ROC curves and LDA coordinates in biplots (**Fig 3E-H**). Label permutation testing confirmed significant discrimination between CADASIL and disease for both cohorts when provided protein level information for proteins in the CADASIL-Late signature (*P* = 0.049 for Late → Early; *P* = 0.017 for Late → Colombia; **Table S3**). The encouraging results of these tests provided further validation for our machine learning approach, indicating that our disease-associated protein set held reliable predictive capacity across diverse CADASIL populations.

### Early and Late CADASIL proteomic signatures have both overlapping and distinct network components suggesting evolution and progression in patho-mechanisms of CADASIL across disease stages

In order to attain a better understanding of the molecular pathways, associated putative mechanisms, and interactions between proteins captured by our protein signatures, we used the web based STRING platform (*15*). Both the Early and Late signatures served as input, and the resulting networks were overlapped to assess similarities and differences. The resulting network revealed 33 nodes and 55 edges, highlighting intricate relationships among proteins (**Fig. 5A**). Several proteins in the signature served as network hubs, including FN1 with 12 interactions, ITGB1 with 8, and SDC4, KNG1, and VEGFR3 each with 7 interactions. Notably, 7 interactions were found between proteins specifically associated with CADASIL-Early and CADASIL-Late, providing insights into potential transitional pathways and the multifactorial nature of CADASIL.

**Fig. 5.**
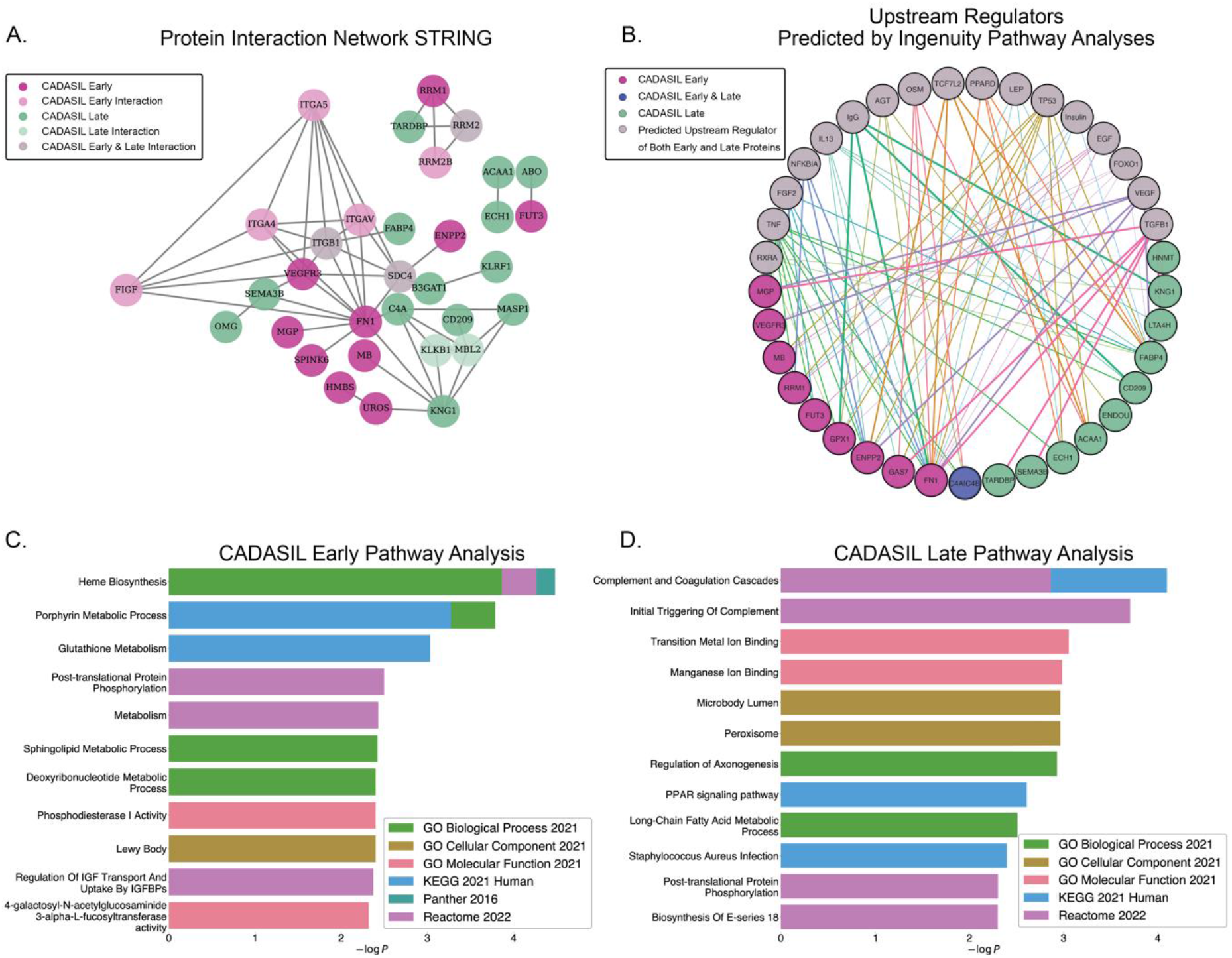
Pathway and network analyses of Early and Late signatures. **(A)** STRING network of Early and Late signature proteins with predicted protein-protein physical and functional interactions. **(B)** IPA network of Early and Late signature proteins as well as predicted upstream regulators. Edges are colored according to the corresponding upstream regulator and edge width was assigned based on predicted activation Z-scores (see also Table S4). **(C)** Collated EnrichR pathway analysis from several libraries using Early signature. **(D)** Collated EnrichR pathway analysis from several libraries using Late signature.

We then made use of Ingenuity Pathway Analysis (IPA) to predict upstream regulators of each protein signature, respectively. 17 proteins were found to be significant upstream regulators of the Early and Late networks (overlap *P* < 0.05). The overlapped regulator network showed notable interconnectedness between CADASIL-Early and CADASIL-Late signatures (**Fig. 5B**). The predicted activation Z-scores of all upstream regulators are presented in **Table S4**. TGF-β1, a protein with hypothesized involvement in CADASIL (*16–18*), emerged as a key regulator protein. FN1, a hub protein in the STRING analyses, also emerged as a hub in the IPA network, sharing many upstream regulators with other CADASIL-Early and CADASIL-Late proteins. The upstream regulator analysis provides potential insights into therapeutic targets and yet to be explored adjacent pathways.

### Specific investigation of signature proteins and their corresponding genes identified links to perturbations in metabolic, neuronal, cell differentiation, and inflammatory pathways

Noting unique components when contrasting the CADASIL-Early with CADASIL-Late signature networks, we then sought to identify specific perturbations linked to the enrichment of pathways or differential expression of proteins. We investigated these potentially pathological associations using both the EnrichR (*19*) platform and a Protein-centric Reverse GEO Search (*20*). For the CADASIL-Early signature (**Fig. 5C**), several enriched pathways were noted. These included: heme biosynthesis (*P* = 3.3 × 10^−5^), specifically the porphyrin metabolism pathway (*P* = 5.3 × 10^−4^), glutathione metabolism (*P* = 9.3 × 10^−4^) and phosphodiesterase activity (*P* = 4.0 × 10^−3^), regulation of axonogenesis (*P* = 1.2 × 10^−3^), and Lewy body enrichment (*P* = 4.0 × 10^−3^). These findings suggest notable changes encompassing metabolic, oxidative stress, and neuronal dysfunction related pathologies. Further, the Early signature Reverse GEO Search (**Figure S1A**) highlighted FN1 as the top hit and linked this association to upregulation in a hepatocellular carcinoma model (*P* = 5.50 × 10^−39^), cytokine-treated insulinoma cells (*P* = 1.29 × 10^−28^) and a proliferating glioblastoma (*P* = 6.31 × 10^−26^).

For the CADASIL-Late signature (**Fig. 5D**), we noted significant enrichment for complement pathways (*P* = 8.0 × 10^−5^), such as the classical complement cascade (*P* = 1.4 × 10^−3^), peroxisome pathways (*P* = 2.5 × 10^−3^), including fatty acid metabolism (*P* = 3.1 × 10^−3^), manganese binding (*P* = 1.0 × 10^−3^), PPAR signaling (*P* = 2.5 × 10^−3^), staphylococcal infection (*P* = 4.0 × 10^−3^), and acetyl-CoA metabolism (*P* = 6.0 × 10^−3^). The broad diversity of enriched pathways observed in this signature reflect key pathologies associated with neuronal dysfunction, chronic inflammation, and metabolic changes. Further, the Late signature Reverse GEO Search (**Figure S1B**) noted significant associations with the upregulation of LTA4H in intrahepatic cholangiocarcinoma (*P* = 2.04 × 10^−30^) and in antifibrotic models (*P* = 8.32 × 10^−27^) as well as elevated ACAA1 in cholangiocarcinoma (*P* = 1.70 × 10^−21^).

### Dynamic transcriptomic changes and cell-specific expression patterns uncovered in CADASIL proteomic signature

To further validate clinical relevance, we compared peripheral blood proteomic signatures against brain tissue bulk transcriptomics signatures, using transcriptomics data generated from 5 CADASIL and 7 control brains from the BA4/6 area of the human cortical region. From the Early signature, HPCAL1 and VEGFR3 showed upregulation in CADASIL patients (log_2_FC = 1.3 and log_2_FC = 0.7, respectively; **Fig. 6A**) while ENPP2, GAS7, and RRM1 were significantly downregulated (log_2_FC = −2.6, log_2_FC = −1.1, and log_2_FC = −0.8, respectively; **Fig. 6A**). From the Late signature, BCAR3 was significantly upregulated in CADASIL (log_2_FC = 1.0, **Fig. 6B**) whereas SEMA3B and OMG displayed significant downregulation in CADASIL brain tissue in comparison to control brain tissue (log_2_FC = −2.2 and log_2_FC = −1.3, respectively; **Fig. 6B**). These results reveal dynamic transcriptomic changes for signature proteins in CADASIL patients compared to controls. It should be noted that all brain tissue represents the end stage.

**Fig. 6.**
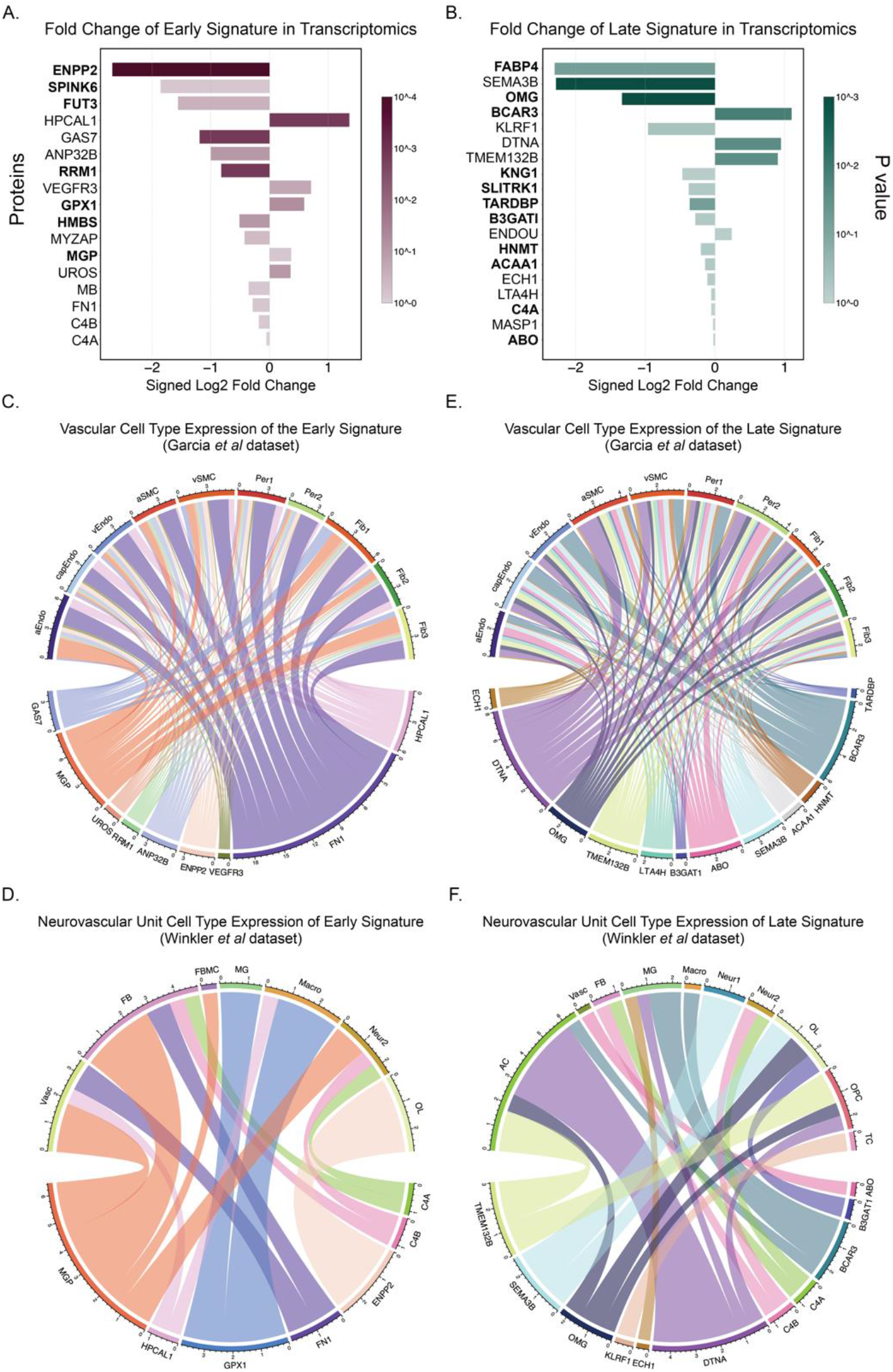
Brain transcriptomic analysis of Early and Late signatures in CADASIL. **(A-B)** Bulk RNASeq analysis depicting fold changes (log2FC) of the **(A)** Early signature and **(B)** Late signature in CADASIL versus control brain tissue. Bolded proteins are co-directional in plasma and brain tissue. **(C-D)** Chord diagrams illustrating vascular cell type expression of the Early signature based on Garcia et al. dataset **(C)**, and Winkler et al. dataset **(D)**. **(E-F)** Chord diagrams showcasing vascular cell type expression of the Late signature based on Garcia et al. dataset **(E)**, and Winkler et al. dataset **(F)**. **(C-F)** The width of the band represents the degree of upregulation of the protein in a specific cell type. **(C, E)** aEndo, arterial endothelial cell; aSMC, arterial smooth muscle cell; capEndo, capillary endothelial cell; Fib1, fibroblast cluster 1; Fib2, fibroblast cluster 2; Fib3, fibroblast clutter 3; Per1, pericyte cluster 1; Per2, pericyte cluster 2; vEndo, venous endothelial cell; vSMC, vascular smooth muscle cell. **(D, F)** AC, astrocyte; Vasc, vascular cells; FB, perivascular fibroblast; FBMC, fibromyocyte; Macro, macrophage; TC, T cell; Neu1, Neuron Cluster 2; Neu1, Neuron Cluster 2; MG, microglia; OL, oligodendrocyte; and OPC, oligodendrocyte precursor cell.

We then analyzed publicly available single-cell RNA-seq data from brain vascular cells to map neurovascular expression of the protein signatures (*21, 22*). These data were generated *ex-vivo* from normal cerebral cortex from patients undergoing surgery for refractory epilepsy and cortical dysplasia. We made use of *ex-vivo* cell atlases, rather than postmortem atlases, due to notable changes in gene expression between living and postmortem brains (*23*).For the Early signature, endothelial expression was prominent for VEGFR3, FN1, MGP, and HPCAL1 (**Fig 5C-D**). ENPP2 was expressed in oligodendrocyte lineage cells. For the CADASIL-Late signature, endothelial expression was noted for SEMA3B, TMEM132B, and ABO (**Fig 5E-F**). OMG and B3GAT1 showed oligodendrocyte expression. KLRF1 was expressed in T cells and BCAR3 in astrocytes. The remainder of the proteins were not significantly cell specific. These cell-specific expression patterns provide insight into which brain vascular cells display dysregulation of signature proteins in CADASIL progression.

### CADASIL signature proteins exhibit strong associations with quantitative clinical measures and serve as predictors of disease-related traits

Lastly, we sought to better understand the clinical relevance of blood proteomic signatures obtained. We began this final investigation by testing the association of our protein signatures with characteristic MRI findings from CADASIL-Early patients. The assessed MRI metrics included white matter hyperintensities (WMH), a measure of white matter injury, enlarged perivascular space volume (ePVS), which are a more specific radiographic measure of small vessel disease(*24*), and brain atrophy measured by the negative log of brain-parenchymal fraction (-log(BPF); with higher −log(BPF) corresponding to higher brain atrophy) (**Fig. 7A-B**). In the Early signature (**Fig. 7A**), ANP32B showed the strongest positive association with WMH (*R* = 0.52, *P =* 0.008) and ePVS (*R* = 0.46, *P* = 0.022). Meanwhile, GAS7 was associated negatively with ePVS (*R* = −0.46, *P* = 0.021) but positively with brain atrophy (*R* = −0.24, *P* > 0.05). For the Late signature (**Fig. 7B**), C4A|C4B was positively associated with measures of ePVS load (*R* = 0.38, *P* > 0.05) and DTNA was positively associated with ePVS (*R* = 0.43, *P* < 0.05). TMEM132B was positively associated with brain atrophy (*R* = −0.35, *P* > 0.05).

**Fig. 7.**
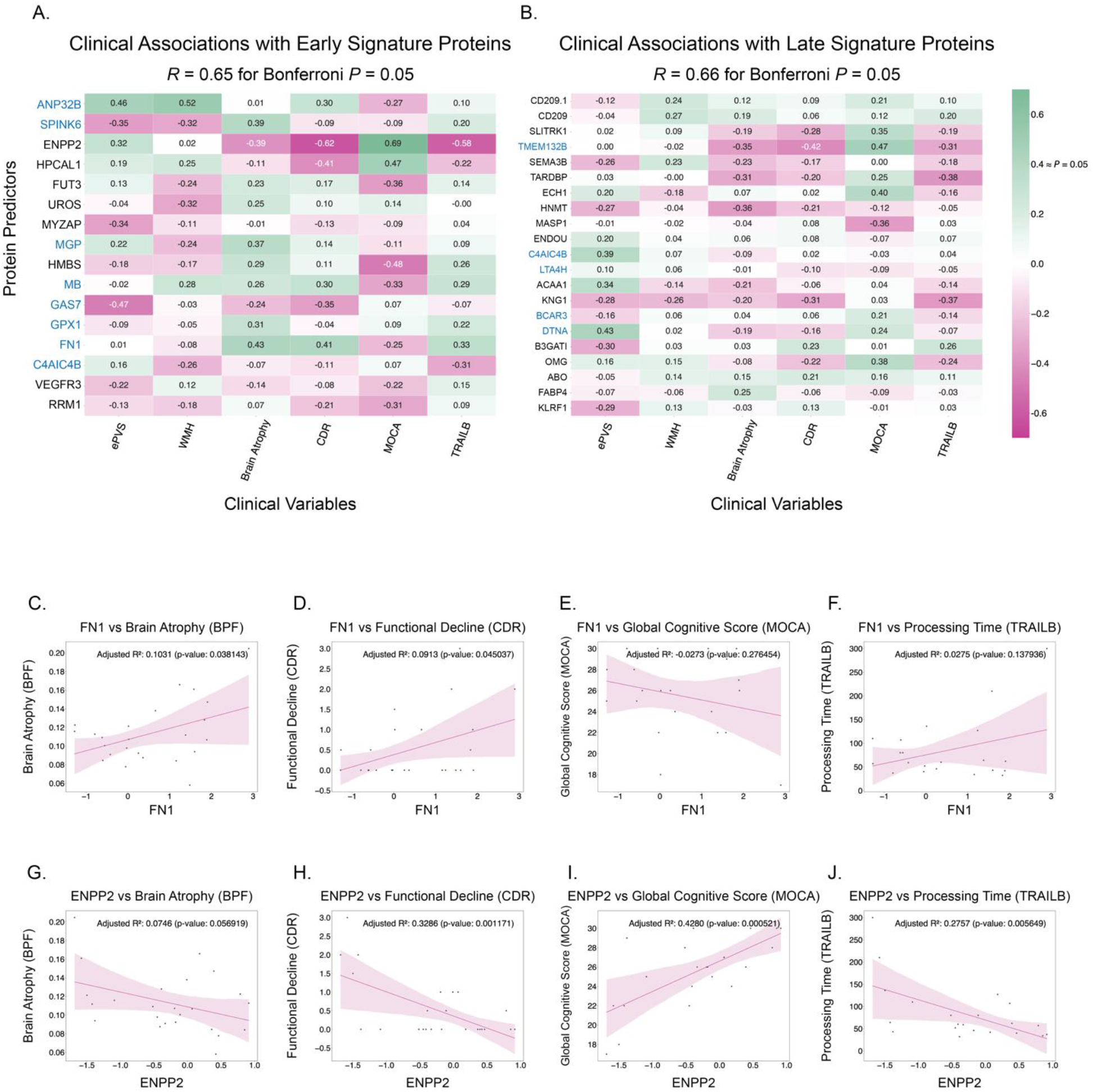
Plasma proteomic signatures in CADASIL patients: association and regression analyses. **(A)** Association analysis of the Early signature. **(B)** Association analysis of the Late signature. Proteins labeled in blue are upregulated in plasma; proteins labeled in black are downregulated in plasma. **(C, D, E, F)** Regression analyses for protein FN1 against various imaging and clinical metrics: **(C)** brain atrophy, **(D)** functional decline, **(E)** global cognition, and **(F)** processing time. **(G, H, I, J)** Regression analysis of protein ENPP2 against various imaging and clinical metrics: (G) brain atrophy (-log(BPF)), **(H)** functional decline (CDR), **(I)** global cognition (MOCA), and **(J)** processing time (TRAILB). Brain Atrophy, BPF, brain parenchymal fraction; Functional Decline, CDR, clinical dementia rating score; ePVS, Enlarged Perivascular Space Volume; Global Cognition, MOCA, Montreal Cognitive Assessment Score; Processing Time, TRAILB, Trail Making Test Part B Completion; Time; WMH, White Matter Hyperintensities. Protein labels colored blue indicate upregulation in CADASIL plasma compared to controls. Proteins colored black indicate downregulation in CADASIL plasma compared to controls.

With regression analyses, we show associations between proteins in the signatures and measures of cognitive impairment in CADASIL-Early patients. The clinical outcomes include functional decline (Clinical Dementia Rating Score (CDR)) (*25, 26*), cognitive processing time (Trail Making Test Part B Completion Time (TRAILB)), and global cognitive score (Montreal Cognitive Assessment score (MOCA)). Due to limited statistical power in clinical data, our regression analyses were underpowered to survive Bonferroni correction. Moreover, given the hypothesis-driven nature of these analyses, we did not think it pertinent to look at multiple comparisons corrected p-values. Instead, our investigation aimed to determine if there was consistency in the associations of proteins across MRI findings and cognitive measures. We sought to understand whether the values of proteins were consistently related across various measures, termed as “congruent” associations. For example, if a protein showed a positive association with increased brain atrophy, clinical congruence would suggest it also showed a positive association with increased functional decline and slower processing speed. Conversely, it would exhibit a negative association with global cognition, as higher global cognition indicates better cognitive outcomes. Thus, we defined a protein as clinically congruent for disease progression if it showed a positive association with brain atrophy, functional decline, and processing speed, and a negative association with global cognition. Conversely, proteins with a potentially disease-ameliorating or neuro-protective role in the neurovascular unit were expected to exhibit the opposite pattern: a negative association with brain atrophy, functional decline, and processing speed, and a positive association with global cognition. FN1, a protein also highlighted in the transcriptomic data, demonstrated clinical congruence with disease progression. FN1 was positively associated with brain atrophy (*R* = 0.42, *P* = 0.038; **Fig. 7C**), functional decline (*R* = 0.41, *P* = 0.045; **Fig. 7D**), and slowed processing speed (*R* = 0.33, *P* > 0.05; **Fig. 7F**). As well, FN1 had a strong negative association with global cognition (*R* = −0.24, *P* > 0.05; **Fig. 7E**). ENPP2, on the other hand, was clinically congruent with a possible disease-meliorating role. ENPP2 was negatively associated with brain atrophy (*R* = −0.39, *P* > 0.05; **Fig. 7G**), functional decline (*R* = −0.62, *P* = 0.001; **Fig. 7H**), and slowed processing time (*R* = −0.58, *P* = 0.006; **Fig. 7I**). Furthermore, ENPP2 was positively associated with global cognition (*R* = 0.69, *P* = 0.0005; **Fig. 7J**).

Other proteins showed associations with cognitive measures. For the CADASIL-Early signature (**Fig. 7A**), HPCAL1 was clinically congruent and negatively associated with functional decline (*R* = −0.41, *P* = 0.043) but positively associated with global cognition (*R* = 0.47, *P* = 0.030). HMBS exhibited a strong negative association with global cognition (*R* = −0.47, *P* = 0.029). For the CADASIL-Late signature (**Fig. 7B**), SLITRK1 showed a consistent negative association with cognition, including higher functional decline (*R* = −0.28, *P* > 0.05) and higher processing time (*R* = −0.18, *P* > 0.05). TMEM132B exhibited positive associations with global cognition (*R* = 0.46, *P* = 0.032).

## Discussion

The goal of our study was to take an unbiased approach to discovery of blood molecular signatures of CADASIL, a monogenic form of VCID, as a critical first step toward development of biomarkers and formulation of mechanistic hypotheses for development of impactful treatments. To this end we formulated a novel experimental design and analytical approach aimed at extracting proteomic signatures in a methodologically unbiased manner, specifically tailored to monogenic diseases or disease states with clear categorization of disease and control groups. This research marks, to our knowledge, the first instance of unveiling a plasma proteomic signature associated with a monogenic form of VCID. Our findings shed light on molecular pathologies in CADASIL and serve as a stepping stone towards a broader investigation of VCID blood signatures.

We undertook an approach for capturing multivariate associations inherent in “-omics” datasets that traditional differential expression analyses reliant on univariate significance cannot capture (*27*). Chowdhury et al. (2023) used a technically analogous approach to probe the proteomics of high-grade ovarian cancer, revealing a highly predictive, externally validated biomarker panel (*28*). Our independently constructed methodology demonstrated a comparably high efficacy. By implementing a heterogenous bagging methodology, we amalgamated multiple algorithms, including recursive feature elimination, linear methods (logistic regression and rLDA), and non-linear methods (random forests and Markov blankets) (*29–33*). This combination mitigates the risk of overfitting to a singular method’s decision boundary, especially given the disparity between our limited sample size and the exceedingly large feature count inherent in proteomic research (*34*). Rigorous cross-validation, permutation testing, and validation through independent cohorts further curtailed the potential for overfitting (*35*). Finally, we used brain tissue transcriptomics and clinical data to ensure that proteins identified by ML in the blood were relevant to the brain and clinical phenotypes of the disease. The advantage of this approach over univariate differential expression analysis encompasses enhanced adaptability in representing nonlinear dynamics and interactions, a reduced propensity for false positives owing to collinearity, a focus on disease categorization beyond mere association, inherent feature selection for pinpointing crucial subsets, and augmented resilience against batch effects through ensemble aggregation (*36, 37*).

Applying this multi-step machine learning workflow resulted in the identification of robust protein signatures in both cohorts. The CADASIL-Early cohort yielded a concise 16-protein signature that primarily illuminated alterations in metabolic pathways. The CADASIL-Late cohort revealed a 20-protein signature characterized by chronic inflammation, immune alterations, and metabolic dysfunction. The CADASIL-Colombia cohort was utilized for external validation as an independently collected cohort. This validation cohort was characterized by large heterogeneity with regard to the age of participants and the age of the samples. Therefore, some technical noise was expected regarding plasma protein levels. Despite this limitation, we externally validated the discriminatory ability of the Late signal (*AUC* = 0.716, *P* < 0.005; **Table S3**), whereas the AUC score of the Early signal was marginally significant (*AUC* = 0.610, *P* = 0.055; **Table S2**). This finding supports our hypothesis that robust proteomic signatures and molecular signals could be identified using our unbiased and multivariate analytical approaches.

Specific investigation of the proteins comprising each signature using a variety of methods, ranging from pathway enrichment analysis to single-cell RNA-seq, highlighted several additional noteworthy observations that may also serve as guides for the development of new therapeutic strategies. Perturbed pathways suggested by the CADASIL-Early signature emphasized disruptions in heme, porphyrin, and glutathione metabolic processes, aligning with pathways such as oxidative stress resistance via glutathione and biosynthesis of porphyrin-containing compounds. CADASIL compromises redox equilibrium and increases the production of reactive oxygen species (*38–40*). This is consistent with the enrichment of the glutathione pathway, which is known for its role in neutralizing oxidative stress.

Interestingly, TDP-43 was identified in the CADASIL-Late signature (TARDP, a multimeric TDP-43 protein). The cytoplasmic mislocalization of TDP-43 and its aggregation are prominent pathologies in many neurodegenerative diseases (Frontotemporal Lobar Degeneration, Amyotrophic Lateral Sclerosis, and Alzheimer’s Disease) (*41, 42*). It is possible that TDP-43 related molecular dysregulations may serve as an explanatory factor for the observed associations between CADASIL and other neurodegenerative diseases (e.g., ALS, FTD) (*43–45*). However, its specific role in CADASIL and its links to *NOTCH3* mutations have yet to be investigated. TDP-43 was found to be downregulated in plasma of CADASIL patients. It is not clear whether it is mis-localized. Similarly, TDP-43 levels were found to be decreased in the plasma of FTD patients (*46*). The presence of TDP-43 in the CADASIL-Late signature suggests a potentially progressive pathology primed for therapeutic targeting, and thus warrants further investigation.

Other hallmark pathologies known to be associated with other neurodegenerative conditions were also highlighted, particularly in pathway analysis of the CADASIL-Early signature. Pathways associated with Lewy bodies, the defining characteristic of Lewy body dementia (*47*) and often observed in Parkinson’s disease (*48*), were found to be significantly enriched (*P* = 4.0 × 10^−3^). This result is intriguing given prior findings that link *NOTCH3* mutations to worsened clinical outcomes in Parkinson’s disease (*49*) and numerous reports observing parkinsonism in late CADASIL (*50*). The finding of Lewy body pathways in CADASIL implicates a possible pathological link between CADASIL and Lewy-body-related neurodegenerative diseases.

TGF-β1 emerged as a primary predicted upstream regulator of both signatures. TGF-β1 signaling has consistently been found to play a functional role in the cerebrovascular system, including vascular senescence (*51–54*), cerebral angiogenesis and maintenance of brain vessel homeostasis (*16*). Furthermore, TGF-β1 and its receptors are known to play a key role in fibrosis across several diseases (*55, 56*). Specific to the pathways affected in our signatures, earlier research has linked TGF-β1 upregulation to heightened oxidative stress and decreased glutathione levels in endothelial cells (*17*). Additionally, latent TGF-β1 has been found to bind mutated NOTCH3 extracellular aggregates, suggesting its potential involvement in fibrogenesis (*18*). The confluence of fibrosis-associated TGF-β1 upstream regulation with the proteomic signature of CADASIL provides biological validation, considering that fibrosis is a characteristic feature of CADASIL histopathology (*57*). Furthermore, Reverse GEO analyses reported connections with many cases of oncological disease, coupled with recent studies emerging from the oncology field to target the TGF-β pathway (*58*), suggest potential analogous alterations in processes such as angiogenesis and present an intriguing path forward for further investigation.

In addition to their relevance in neurodegenerative and fibrotic pathways, the Early and Late signatures in our study also revealed associations with past proteomic analyses related to aging and cerebrovascular dysfunction. Oh et al. employed a machine learning-based approach to create a Feature Importance for Biological Aging score for human blood plasma proteins measured on the same SomaLogic platform as our study (*59*). Their investigation into over 4,000 proteins pinpointed a distinct five protein signature of aging arteries, notably including MGP, a protein also identified in our Early signature. Significantly, MGP showed a substantial interaction with TAGLN in their analysis, a protein under the regulatory influence of TGF-β1 (*60*) and the most significant protein in the Oh et al. organismal aging model. The interaction of MGP and TAGLN, both prominently expressed in fibroblasts and endothelial cells, underscores their potential role in the advanced vascular aging process characteristic of CADASIL, and possibly in the general aging population. Similarly, Walker et al. found that plasma TAGLN is highly significant for an increased risk of dementia, further underscoring the relevance of MGP/TAGLN proteins in aging-related pathologies (*61*). Intriguingly, the adipose aging signature of Oh et al., which demonstrated the third highest hazard ratio for organ-chronological age-gap, included FABP4, a protein of the Late signature. The co-presence of these signature proteins in our age-matched cohorts is particularly noteworthy. It may characterize CADASIL as an expedited model for studying cerebrovascular aging, extending beyond its primary neurodegenerative context. This aspect of our findings not only reinforces their robustness but also enhances the generalizability and applicability of our results in broader aging research.

We further validated our peripheral blood signature proteins against brain tissue transcriptomics and assigned proteins to specific cells using publicly available single nucleus RNA-seq data. Using the bulk transcriptomic data generated, we found dynamic transcriptomic changes. Additionally, using *ex-vivo* single-cell RNA-seq data from brain vascular cells (*21, 22*), we observed cell-specific expression patterns consistent with pathological alterations in the cerebrovasculature. Endothelial cells and fibroblasts appeared to be prominently affected in the Early signature, while the involvement of astrocytes, microglia, and oligodendrocytes was noted in the Late signature, consistent with the advancement of disease from brain borders into parenchyma as disease progresses. This suggests that therapeutic targets may differ based on the disease stage. Both signatures showed marked expression of numerous proteins (FN1, MGP, GAS7, DTNA, and TMEM132B) in fibroblasts, perhaps further reflecting fibrosis, collagen protein alterations, and extracellular matrix changes identified by pathway analyses. Numerous of these molecular processes and pathways have been implicated in CADASIL, including collagen/ECM involvement (*4, 62–64*) and fibrosis (*17, 40*).

Lastly, to examine the clinical relevance of our molecular findings we tested associations between identified proteomic signatures and disease-associated quantitative clinical phenotypes. For this we limited our analysis to CADASIL-Early patient data. We found strong associations with clinical congruence across several metrics. Several proteins from the CADASIL-Early signature were found to be associated with neuroimaging findings. Fibronectin, a protein previously known to be enriched in CADASIL vessels and shown to increase levels in blood vessels of CADASIL patients (*18*), was upregulated and demonstrated clinical congruence with disease progression in the CADASIL-Early cohort. While FN1 plays a critical role in vascular remodeling after hypoxia- or hypertension-induced vessel injury (*65, 66*), its aggregation in tissues may be detrimental by promotion of thrombogenesis (*67, 68*), neuroinflammation (*69*), and arrest of myelination (*70*), all thought to be components of chronic cSVD and stroke-related VCID. Upregulated CADASIL FN1 levels were positively associated with increased cerebral atrophy and cognitive deterioration in our Early cohort, emerging as a critical, possibly neurodegenerative factor in CADASIL, which warrants exploration in future studies. Conversely, ENPP2 (autotaxin) displayed clinical congruence with a disease-meliorating character, as it was inversely associated with brain atrophy and cognitive decline measures. ENPP2 was significantly downregulated in both plasma and brain transcriptomics of CADASIL patients. ENPP2 is an abundantly expressed member of the ectonucleotide pyrophosphatase/phosphodiesterase family with lysophospholipase activity, catalyzing lysophosphatidic acid (LPA) formation (*71*). Despite mounting evidence for an excitotoxic potential in acute stroke (*72*), other studies have noted potentially beneficial effects in oligodendrocyte maturation (*73*), protection of endothelial cells from hypoxia (*74*) and suppression of CD8+ T cell infiltration in tumors (*75*). These results suggest that ENP22 might confer protection in chronic, but not acute injury, and thus its depletion could reflect long-term changes in CADASIL rather than the effects of strokes.

The elucidation of distinct Early and Late signatures that discriminate CADASIL from controls and are associated with clinical outcome metrics represents an advancement in the development of molecular approaches to advance precision diagnostics across disease stages. We compensated for small sample sizes by using novel analytical methods in addition to cross-validation of results in independent cohorts and brain tissue. The methods for integration of advanced computational techniques and independent validation cohorts translate cross-sectional discoveries into robust and generalizable targets. These findings lay the groundwork for the elucidation of CADASIL pathogenesis and the subsequent identification of disease-stage-specific biomarkers and disease-stage agnostic or specific therapeutic targets.

To our knowledge, this is the first study to implement a multi-step machine learning approach to blood proteomics data to uncover molecular signatures as an unbiased starting point for understanding evolving molecular dysregulation across disease stages in CADASIL. The application of multidimensional analytical techniques enabled us to capture interactions between proteins and patterns shared among multiple analytes in a single analysis, rather than limiting our investigation to specific hypothesis testing in instances of univariate significance. This approach can provide a more comprehensive view of the dataset than most proteomic studies currently in the literature by directly interrogating interactions in a high-dimensional space. The broader adoption of this approach or similar multidimensional pipelines may provide the opportunity to gain a more complete understanding of “-omics” datasets and guide future research towards novel therapeutic targets.

## Materials & Methods

### Study Design - Overview of cohorts

Our study employed plasma proteomics data and clinical information from diverse cohorts, described below. Information regarding demographics (age, sex) and neurological function were provided. Study protocols were approved by their respective Institutional Review Boards. Research was performed in accordance with the Code of Ethics of the World Medical Association. Written informed consent was obtained from all patients before data collection.

### Early CADASIL cohort (*N* = 53)

CADASIL patients (*N* = 25) were consecutively recruited, so as to avoid bias, at the Memory and Aging Center, UCSF, between February 25, 2019, to August 2, 2021. Patients were evaluated by neuropsychological testing, subjected to a blood draw (for plasma collection), and, except for one, underwent MRI neuroimaging. Neurocognitive testing included measures of functional decline (Global Clinical Dementia Rating - CDRTot, and Sum of Boxes - CDRBox), global cognition (Montreal Cognitive Assessment - MOCA), and processing speed (Modified Trail Making Test completion time - MTTime, Trail Making Test B completion time - TRAILB). MRI-derived measurements included white matter hyperintensity (WMH) volume, enlarged perivascular space (ePVS) volume (measured by LOAD), and Brain Parenchymal Fraction (BPF), which were quantified according to a previously described image processing pipeline (*76*). CADASIL status was confirmed based on *NOTCH3* sequencing. The inclusion criteria for all control subjects (*N* = 28) were intact daily functioning per an informant (Clinical Dementia Rating = 0), neuropsychological performances within normative standards, and absence of significant clinical neurological disease assessed by history and physical exam. Control subjects underwent blood collection but no neuropsychological evaluation or medical imaging.

### Mayo Clinic CADASIL cohort (*N* = 45)

CADASIL patients (*N* = 20) were recruited at the Department of Neurology, Mayo Clinic, Jacksonville between July 29, 2014, to February 2, 2021. Patients underwent clinical evaluation and blood draw. CADASIL status was confirmed based on *NOTCH3* mutations discovered via genetic sequencing. Control subjects (*N* = 25) were selected based on absence of significant clinical neurological disease assessed by history and physical exam.

### Colombia CADASIL cohort (*N* = 66)

Patients with CADASIL from Colombia (N = 25) and controls with no *NOTCH3* mutations (*N* = 41) who were family members of the CADASIL patients, were recruited from a cohort in Colombia during two periods, from August 3, 2000, to July 14, 2005 and from January 12, 2015 to December 11, 2016. A subset of participants (10 CADASIL, 20 control participants) was longitudinally evaluated during both periods and constituted the longitudinal cohort for this study. Sequencing the *NOTCH3* gene confirmed CADASIL status.

### Plasma Collection and Proteomic Analysis

For proteomics characterization, plasma samples from all cohorts were analyzed through the SOMAscan 7k assay (SomaLogic, Inc., Boulder, CO). The SomaScan assay offers the advantage of unbiased protein expression analysis of a wide range of proteins, covering all biological functional domains. As described previously, the SOMAscan assay kit employs highly selective single-stranded modified Slow Off-rate Modified DNA Aptamers (SOMAmer) for protein identification and quantification. A custom DNA microarray (Agilent) was used for quantification, which is reported as relative fluorescence units (RFU). Raw data then underwent quality control, calibration, and normalization. Prior to data analysis, we performed sample pre-processing. All non-human SOMAmers (307 proteins) were removed from the dataset leaving 7,289 proteins of the 7,596 proteins measured by SomaLogic. The data was then transformed by a natural log.

### Machine Learning Pipeline

#### Recursive Feature Elimination, Leave-One-Out-Cross-Validation: Definition and Analysis

In our study, we aimed to uncover CADASIL disease-associated changes in proteomic networks using comprehensive and unbiased machine learning (ML) techniques. Our approach’s central basis was that protein sets which are crucial in distinguishing disease states may be key biological drivers of the disease (*32*). We developed a novel ML methodology that employs auxiliary Markov blanket feature selection (*77, 78*) combined with multiple recursive feature selection algorithms to mitigate bias towards any specific algorithm (*79*) and reduce overfitting, which is the fundamental challenge considering the inherent low sample size and high dimensionality of our, and many others, proteomics datasets. The first step of our method was the creation of Leave-One-Out (LOO) partitions of our data (*35*). For a dataset with N samples, we generated N partitions (LOO folds), each excluding one unique sample while including the rest. This approach ensured that the influence of any single sample is minimized in the model, addressing overfitting, and improving the model’s generalizability.

Before feature selection, every protein in each LOO fold was subjected to a univariate t-test at an alpha level of 0.05. This step is vital to reduce the risk of including proteins that show apparent but spurious associations with the disease due to random variation, a common issue in datasets where *n* ≪ *p*. For each fold, ~1,300 proteins survived the filtering step and proceeded to feature selection.

For feature selection, we utilized a diverse array of algorithms, some employing Recursive Feature Elimination (RFE) and others independent of it. RFE is a technique that systematically removes the least significant features to identify the most relevant ones. In the context of our high-dimensional data, RFE helps in reducing the feature space, making the model more robust and less prone to overfitting. The RFE algorithm was coupled with repeated 10-fold cross-validation during each elimination step to minimize variance in selection. The suite of algorithms employed included RFE with Logistic Regression (LR) with L1 and L2 regularization penalties, respectively (*30, 31*), RFE with regularized Linear Discriminant Analysis (rLDA) (*80*), RFE with Random Forests (RF) (*29*), Boruta - Random Forests (*81*), and Maximum-Relevance-Minimum-Redundancy (MRMR) with an F-Statistic evaluator (*82*). Markov blanket feature selection was employed separately on the original datasets, due to computational expense and subsequently incorporated during the later aggregation steps (*77, 78*).

Each algorithm was utilized to address specific challenges in the *n ≪ p* problem, where variables significantly outnumber observations. RFE-LR with L1 regularization was employed for its sparsity-inducing property, efficiently eliminating less significant features in well-defined classes (*31*). RFE-LR with L2 regularization was used to manage multicollinearity, shrinking coefficients without excluding any features, thus preserving the contributions of all variables even in the presence of high inter-correlations (*83*). Regularized-LDA (rLDA) was selected for its enhanced accuracy in settings with normally distributed within-class proteins and small sample sizes, addressing the instability issues of logistic regression in scenarios with significant class separation (*80*). Additionally, the Boruta-RF method was integrated to effectively identify crucial features in high-dimensional datasets by comparing real features against randomly generated shadow features, optimizing feature selection under the n <<< p constraint. The Markov Blanket approach was effective for focusing on complex, non-linear variables most relevant to the target, thus efficiently reducing dimensionality (*77*). Finally, the MRMR method was used for balancing feature relevance and minimizing redundancy, crucial for predictive accuracy in datasets with numerous features. This comprehensive approach demonstrates a nuanced handling of feature selection in complex, high-dimensional datasets.

Each aforementioned algorithm was applied to all *N* LOO folds, producing *N* sets of proteins for each algorithm. The final step in our methodology was a consensus aggregated feature selection approach to combine these results. First, we identified a ‘bag of features’ for each algorithm, selecting only those proteins that appeared in every set produced by that algorithm across all LOO partitions. Next, we cross-referenced these bags of features across different algorithms, further minimizing bias towards any single ML approach. A protein was included in our final proteomic signature only if it appeared in the ‘bag of features’ sets of at least two different models, often appearing in several. This method ensures a consensus among different algorithms, further reducing the risk of model overfitting and bias towards any single ML approach and providing a more reliable indicator of disease-associated proteomic changes.

Subsequently, we performed Principal Component Analysis (PCA) and Linear Discriminant Analysis (LDA) on this protein set for visualization of disease discrimination. PCA was executed to display the remaining variance in the data, emphasizing that it is based on the disease state (*84*). In contrast, LDA was utilized to visually demonstrate that the protein set harbors information crucial for delineating and clustering the disease. Lastly, we performed permutation analysis to evaluate the significance of our findings. We randomly selected a protein signature of the same length from the dataset and evaluated its performance. Similarly, we permuted the class labels and evaluated the resulting classifier performance to ensure the signature was disease specific.

#### Cross-Validation Between Cohorts and External Validation with Independent Cohort

Following the determination of our protein signatures, we proceeded to evaluate their capacity to classify diseases. This exploration was underpinned by the hypothesis that proteins integral to successful disease classification models could potentially serve as disease-associated markers. To this end, we scrutinized the efficacy of an array of machine learning algorithms, inclusive of Linear Support Vector Machines (SVM), Random Forests (RF), Regularized Linear Discriminant Analysis (rLDA), Logistic Regression (LR), Ridge Classifier, Perceptron and Decision Trees. To ensure robustness and applicability of our findings, we sought to validate our proteomic signature specific to CADASIL. This validation was undertaken by refitting the Early disease signature to the Late disease cohort and vice versa. As well, an external CADASIL dataset (CADASIL-Colombia) was tested. This cross-validation and external validation served to substantiate the generalizability of our CADASIL proteomic signatures across different patient cohorts.

### Programming Languages and Packages Used

Data processing was handled in the R environment version 4.2.1 (R Foundation for Statistical Computing, 2022). The more complex machine learning data analyses, including Boruta-Random Forests and MRMR feature selection, were performed using Jupyter Notebooks, leveraging pertinent libraries such as SciKit-Learn for machine learning algorithms and MRMR_Selection for MRMR analyses. SciKit-Learn was also the tool of choice for performing all correlation calculations. For data organization and basic numerical operations, we employed the Pandas and Numpy libraries respectively. Finally, visualizations pertaining to Principal Component Analysis (PCA) and Linear Discriminant Analysis (LDA) were generated using Matplotlib.

### Pathway and Gene Ontology Analysis

Biological function analysis and pathway analyses via the Gene Ontology, KEGG, Panther, and the Reactome databases were performed through the use of an Appyter-based version of the EnrichR web tool (*19*). Similar EnrichR terms from separate databases were combined under one term and depicted as overlapping plots. We used Ingenuity® Pathway Analysis (QIAGEN, Redwood City) software for functional pathway and upstream regulatory analysis (URA) of the proteins–of-interest identified in this study (*85*). For the above we set the significance level threshold for the Benjamin-Hochberg adjusted p-value at 0.05. We further explored proteins of interest in the STRING database version 12.0 for protein-protein functional and physical interaction analysis, the results of which were displayed as a functional network. Interactions were considered with a medium confidence score of 0.4 or higher. We used the Reverse GEO Search Appyter to investigate the disease relevance of the protein signature (*20*). We searched the results for each protein in the signature and plotted the top 25 terms based on the negative log of the p-value for upregulated and downregulated signatures respectively.

### Human Brain Tissue and Bulk Transcriptome Sequencing

We performed RNA sequencing (RNA-seq) analysis on postmortem frontal cortex samples from CADASIL patients (*N* = 5, provided by the Wang Laboratory, University of Michigan) and controls (*N* = 7, provided by the Mount Sinai Neuropathology Brain Bank). In short, deep-frozen samples from the BA4/6 area of the human cortex were micro-dissected. RNA extraction, preparation of cDNA libraries and transcriptome sequencing using the Illumina NovaSeq (paired-end 150-nucleotide read length) was conducted by Novogene Co., LTD (Beijing, China). All samples were assessed to have an RNA integrity number (RIN) above 3.5, so as to ensure a sufficient relative abundance of full-length transcripts. Reads of low quality or containing adapter sequences were filtered. Raw fastq files were analyzed through an in-house transcriptomics pipeline, RAPiD-nf, implemented in the NextFlow framework. Briefly, remaining adapter sequences were filtered by Trimmomatic v0.36 (*86*), STAR v2.7.1 (*87*) was used to align to the hg38 build of the human reference genome (GRCh38), and featureCounts performed BAM-level quantification (*88*). Results were subjected to quality control using FastQC and Picard. For Differential Expression Analysis we employed the DESeq2 package in R (*89*). Before analysis, sequences with sum of sequence counts < 10 across all participants were removed. We calculated the average log2 fold change for each protein between CADASIL and control groups and then plotted the results in a bar plot to visualize transcriptional alterations of the signature proteins.

### Investigation of Brain Vasculature Expression of CADASIL Plasma Signature

To elucidate cell type–specific dysregulation of the CADASIL signature proteins, we analyzed published single-cell RNA-seq data from Winkler et al. and Garcia et al. Heatmaps were generated to visualize the signed log2 fold changes of signature proteins across different brain vascular cell types.

### Statistical Analysis

Mean demographics, imaging, and cognitive test measures were compared between cohorts with Student’s t-test for continuous variables and Chi-square tests for categorical variables. Average signed log2 fold change and Student’s t-test was calculated for old versus new time point CADASIL and control samples in longitudinal Colombia cohort. Linear regression models, with proteins derived from the ML pipeline as predictors, were used to test for associations with imaging and clinical variables. As well, we developed multiple linear regression models with backward feature selection of the protein set to generate feature subsets to predict clinical variables. All statistical analyses were performed using Numpy, Scipy, and Pandas libraries in Python. All statistical tests were unpaired. A 2-sided p-value ≤ 0.05 was considered statistically significant, and a p-value < 0.10 but greater than 0.05 marginally significant.

## Acknowledgments

This work was funded by the National Institute on Aging and Department of Veterans Affairs (IK2CX002180), Rainwater Charitable Foundation (2790954), and Chan Zuckerberg Initiative (2022-316712) (FME). High power computing at the Icahn School of Medicine at Mount Sinai is supported in part through the computational and data resources and staff expertise provided by Scientific Computing and Data and the Clinical and Translational Science Awards (CTSA) grant UL1TR004419.

## Author Contributions

JNK, HR, NK, and FME developed the concept, planned and analyzed data. JNK, HR, and NK performed the biostatistical analyses and generated figures. JNK and FME drafted, reviewed, and edited the final version of the manuscript. MMW provided brain tissue and SF and LM extracted protein from brain tissue. JFM shared plasma samples and JFAV, YTQ and FGL shared data. All authors critically read and provided edits. FME supervised the project, provided the resources, and acquired funding.

## Data and materials availability

All data associated with this study are presented in the article and or the Supplementary Materials. All de-identified patient data is presented in the manuscript and will be shared upon request.

## Supplementary Material

**Fig. S1.**
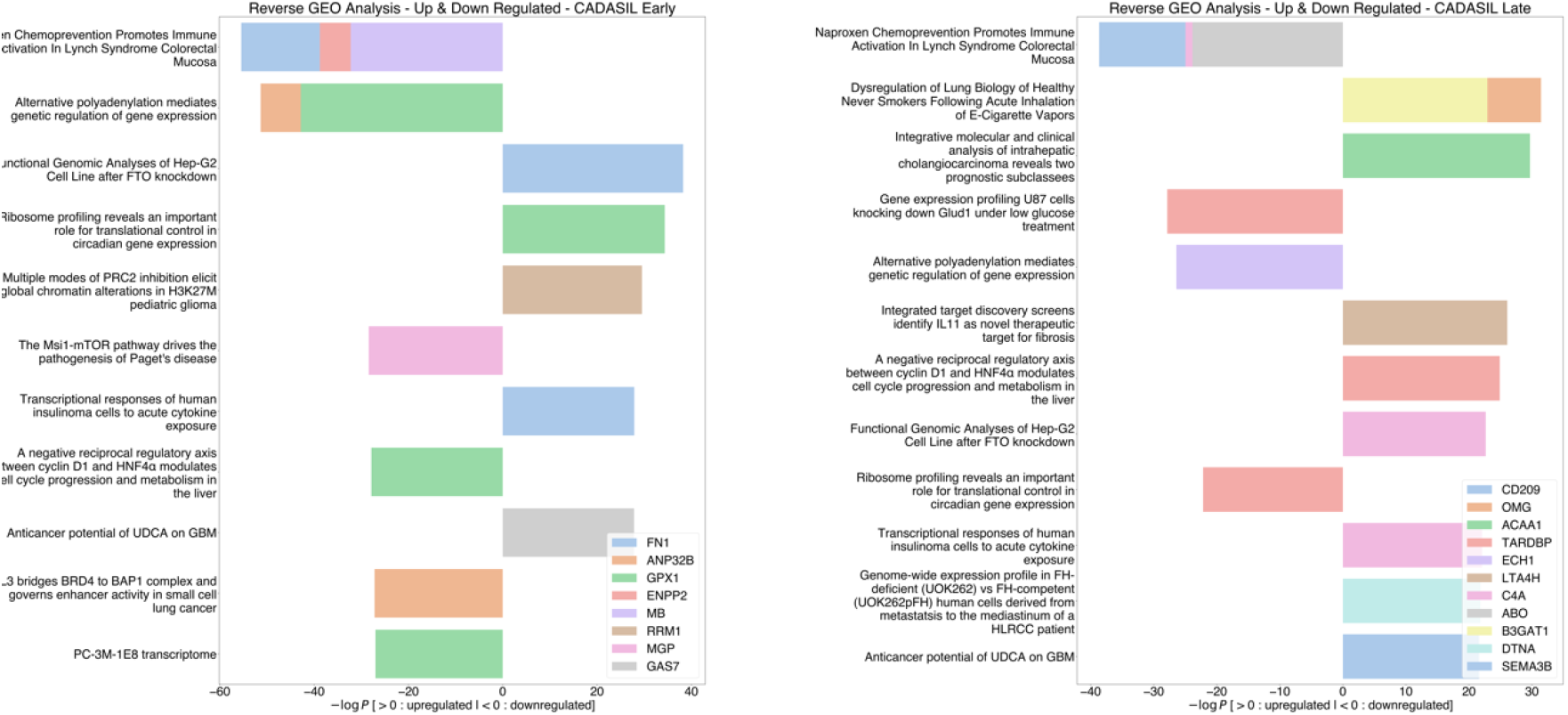
Protein-centric Reverse GEO Search of **(A)** CADASIL-Early and **(B)** CADASIL-Late Signature.

**Table S1.**
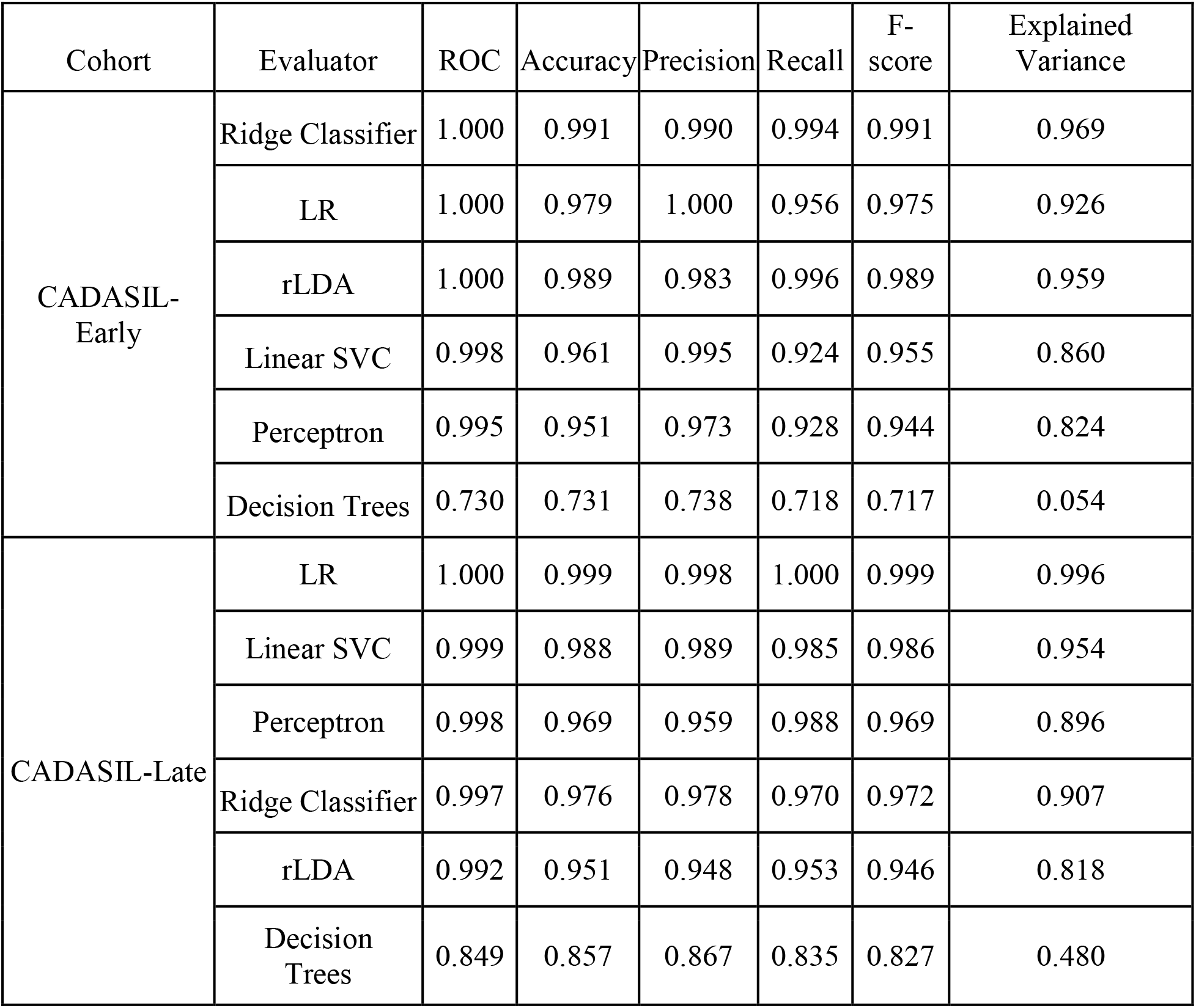
CADASIL-Early ML classifiers and their performance using the CADASIL-Early, CADASIL-Late, and CADASIL-Colombia protein signature.

| Cohort | Evaluator | ROC | Accuracy | Precision | Recall | F-score | Explained Variance |
| --- | --- | --- | --- | --- | --- | --- | --- |
| CADASIL-Early | Ridge Classifier | 1.000 | 0.991 | 0.990 | 0.994 | 0.991 | 0.969 |
|  | LR | 1.000 | 0.979 | 1.000 | 0.956 | 0.975 | 0.926 |
|  | rLDA | 1.000 | 0.989 | 0.983 | 0.996 | 0.989 | 0.959 |
|  | Linear SVC | 0.998 | 0.961 | 0.995 | 0.924 | 0.955 | 0.860 |
|  | Perceptron | 0.995 | 0.951 | 0.973 | 0.928 | 0.944 | 0.824 |
|  | Decision Trees | 0.730 | 0.731 | 0.738 | 0.718 | 0.717 | 0.054 |
| CADASIL-Late | LR | 1.000 | 0.999 | 0.998 | 1.000 | 0.999 | 0.996 |
|  | Linear SVC | 0.999 | 0.988 | 0.989 | 0.985 | 0.986 | 0.954 |
|  | Perceptron | 0.998 | 0.969 | 0.959 | 0.988 | 0.969 | 0.896 |
|  | Ridge Classifier | 0.997 | 0.976 | 0.978 | 0.970 | 0.972 | 0.907 |
|  | rLDA | 0.992 | 0.951 | 0.948 | 0.953 | 0.946 | 0.818 |
|  | Decision Trees | 0.849 | 0.857 | 0.867 | 0.835 | 0.827 | 0.480 |

**Table S2.** Best ML classifiers performance metrics using the CADASIL-Early protein signature on different cohorts.

| Cohort<br>Best ML Model<br>ROC |  |  | *P* value of ROC |  |
| --- | --- | --- | --- | --- |
|  |  |  | Based on ROC Score Distribution of<br>Random Protein Predictor Models | Based on ROC Score Distribution of<br>Permuted Label Models |
| CADASIL-Late | LDA | 0.575 | 0.2058 | 0.3312 |
| CADASIL-Colombia | SVC | 0.610 | 0.0552 | 0.1308 |

**Table S3.** Best ML classifiers performance metrics using the CADASIL-Late protein signature on different cohorts.

| Cohort<br>Best ML Model<br>ROC |  |  | *P* value of ROC |  |
| --- | --- | --- | --- | --- |
|  |  |  | Based on ROC Score Distribution of<br>Random Protein Predictor Models | Based on ROC Score Distribution of<br>Permuted Label Models |
| CADASIL-Early | SVC | 0.765 | 0.0042 | 0.049 |
| CADASIL-Colombia | SVC | 0.716 | 0.0066 | 0.0168 |

**Table S4.**
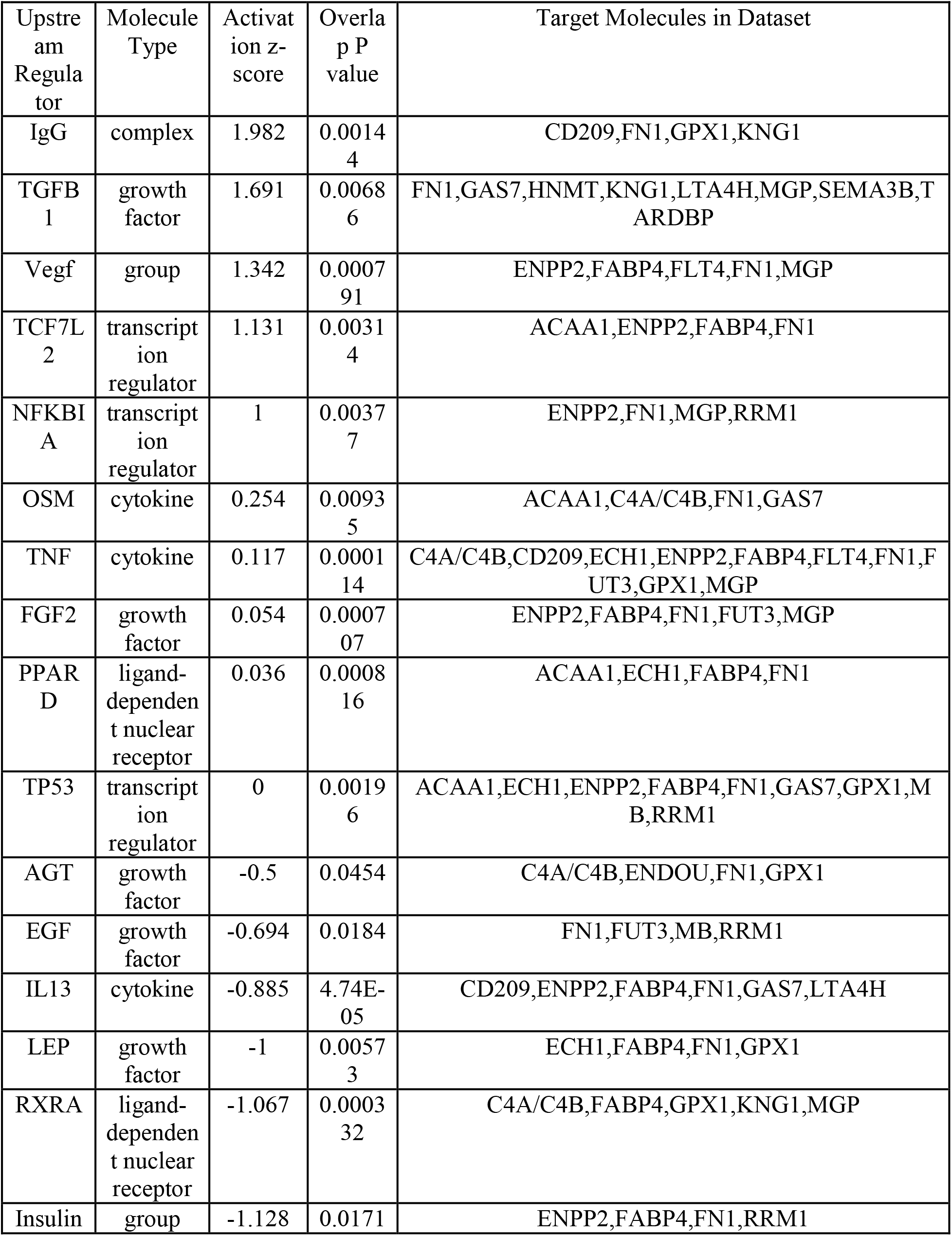
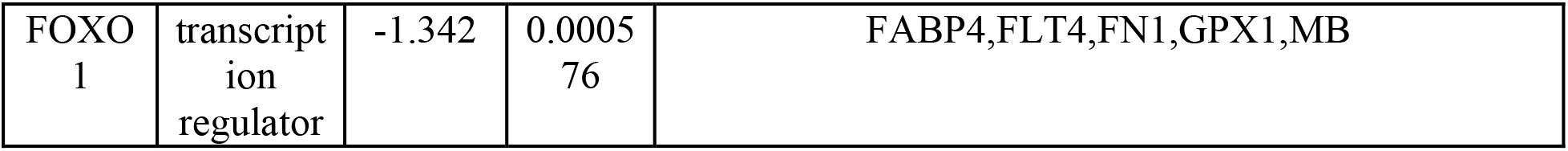
IPA Predicted Upstream Regulators of Early and Late signature proteins.

**Table S5.**
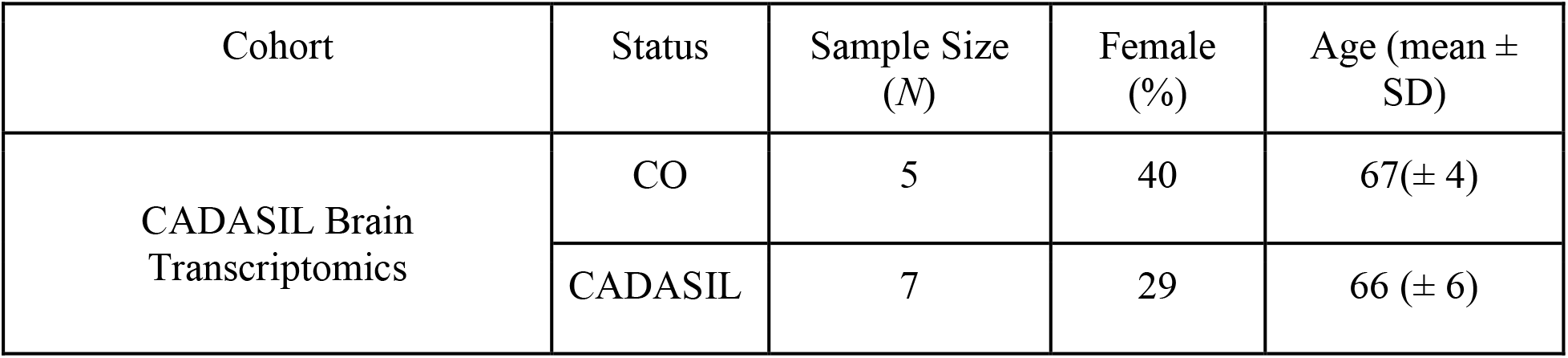
Summary Demographic Data of CADASIL Brain Tissue Transcriptomics Cohort. CO, healthy control; CADASIL, CADASIL cases.

